# Asymmetric detection of relative speed during motion-based figure-ground segregation

**DOI:** 10.64898/2026.11.04.764403

**Authors:** Ying Cao, Bryce Arseneau, Xin Huang

## Abstract

Visual motion can support figure-ground segregation when a local region moves at a different speed from its background, but whether faster- and slower-than-background targets are detected equally well, and how any asymmetry depends on background speed and motion reliability remains incompletely understood. We measured relative-speed detection thresholds in four human observers using an eight-alternative forced-choice localization task. On each trial, one of eight random-dot patches at 10° eccentricity moved either faster or slower than a surrounding moving background. Relative-speed detection thresholds were measured with adaptive staircases at five background speeds (2.5–30°/s) under 100% and 60% motion coherence. In every observer, slower targets required larger relative-speed differences for detection than faster targets. This increment-decrement asymmetry was strongest at low background speeds and diminished as speed increased. Reducing motion coherence elevated thresholds overall but disproportionately impaired detection of slower targets, amplifying the asymmetry, most strongly at the lowest measurable speeds. Under 100% coherence, increment detection showed approximately Weber-like scaling from 10 to 30°/s, whereas under reduced coherence both increment and decrement Weber fractions decreased with increasing background speed. Relative-speed detection therefore depends on the sign of the speed difference, background speed, and motion reliability. The faster-target advantage is consistent with faster-speed weighting reported in cortical area MT and with natural-scene statistics showing that figural regions tend to move faster than their backgrounds, suggesting that relative-speed sensitivity may contribute to motion-based figure-ground segregation. More broadly, such increment-decrement asymmetries can reveal biases in how sensory signals are represented and read out.

## Introduction

Figure-ground segregation allows the visual system to distinguish objects from their background. Motion is a powerful cue for this computation: a region moving differently from its background can be perceived as a distinct figure even when luminance, color, or texture cues are weak or absent (Braddick, 1993; Regan et al., 1992). Suppressive and antagonistic center–surround interactions in motion-sensitive cortex can emphasize motion contrast and contribute to figure– ground segregation (Allman, 1985; Born & Tootell, 1992; Born, 2000; Tadin et al., 2003; Huang et al., 2007, 2008). However, figure and ground signals are not always separated into the center and surround of a receptive field (RF). A motion-defined border may pass through the classical RF; spatially separated figure and ground regions may co-occupy a larger RF; and overlapping or partially occluding surfaces may reside in the same region while moving at different speeds, directions, or depths. In these situations, segregation depends on how the visual cortex represents and weights multiple motion signals within a shared spatial field.

Area MT is well-suited for this computation. MT neurons are selective for motion direction, speed, and binocular disparity, and their relatively large RFs allow them to pool signals across local image regions (Maunsell & Van Essen, 1983; Albright, 1984; DeAngelis et al., 1998; Born & Bradley, 2005). MT also plays a causal role in motion perception (Newsome & Paré, 1988; Salzman et al., 1990, 1992; Rudolph & Pasternak, 1999). Critically, MT responses are not limited to representing a single motion vector: neurophysiological studies using transparent and spatially separated moving stimuli show that MT neurons can represent multiple moving components within their RFs (Snowden et al., 1991; Qian & Andersen, 1994; Recanzone et al., 1997; Krekelberg & van Wezel, 2013; Xiao & Huang, 2015; Chakrala et al., 2024; Wiesner et al., 2020, 2026). Because MT RFs can encompass multiple motion signals, figure-ground segregation may depend not only on center-surround antagonism but also on competition among motion signals represented within the same RF.

Speed is one dimension along which such competition may be organized. MT neurons are tuned for stimulus speed on an approximately logarithmic scale (Maunsell & Van Essen, 1983; Nover et al., 2005), so that equal ratios of speed, rather than equal absolute differences, correspond to comparable distances in the neural representation. This provides a neural account of Weber-like speed discrimination, in which just-detectable speed differences scale with the reference speed (Orban et al., 1985; De Bruyn & Orban, 1988; Nover et al., 2005; Zhang & Stocker, 2022). Most studies of speed discrimination, however, have treated two speeds symmetrically, asking which of two stimuli is faster or whether a single stimulus changed speed. Relatively few studies have directly compared detection of targets faster and slower than the background during motion-based segregation (Sekuler, 1990; Ivry & Cohen, 1992). There are reasons to expect that relative-speed detection is not sign-symmetric. Natural-scene statistics show that figure and ground motion are not equivalent: regions corresponding to figures tend to move faster than the surrounding ground (Friedman et al., 2025). A stimulus moving faster than its background is more consistent with the natural statistical relationship between figure and ground than one moving more slowly. If the visual system is adapted to figure-ground segregation and this statistical regularity, relative-speed detection may be biased toward faster stimuli.

A recent neurophysiological study provides a specific perceptual prediction. When two speeds are present simultaneously within the RF of MT neurons, the population response is biased toward the faster speed, and this bias is strongest at relatively low speeds (Huang et al., 2026). If the representation used for figure-ground segregation based on relative speeds inherits this weighting, a target moving faster than its background should be easier to detect than one moving slower, and the resulting perceptual asymmetry should also be strongest at low background speeds.

Related perceptual asymmetries have been reported for luminance, motion coherence, and temporal speed changes (Whittle, 1986; Wei et al., 2023; Hohnsbein & Mateeff, 2002). The most direct precedent is Sekuler (1990), who measured localization thresholds for speed-defined regions across a wide range of base speeds. Under easier unidirectional conditions, increment and decrement thresholds were similar; when the task was made more difficult by introducing oppositely moving dots or motion noise, a speed-dependent asymmetry emerged, with an increment advantage at low speeds and a decrement advantage at high speeds. Sekuler attributed this crossover to a sigmoidal speed-response function that was accelerating at low speeds and compressive at high speeds. These findings established that motion segregation can depend on the sign of a speed difference. However, the study was conducted near the fovea, and most of the speed range was characterized extensively in a single observer. It also remains unclear whether such asymmetries reflect nonlinear encoding of individual speeds or the way multiple speed signals are weighted when represented within a common receptive field, as in area MT. Here, we measured detection thresholds for targets moving faster and slower than a surrounding random-dot background across five background speeds and two coherence levels. At the eccentricity and spatial scale tested, target and adjacent background motion could plausibly co-occupy the same MT RFs (Albright & Desimone, 1987; Raiguel et al., 1995). Guided by natural-scene statistics and faster-speed weighting in MT, we predicted lower thresholds for faster targets, especially at low speeds. Consistent with this prediction, we found that increment thresholds were lower than decrement thresholds, the asymmetry was strongest at low speeds, and reduced coherence amplified it by disproportionately impairing detection of slower targets.

## Methods

### Subjects

Four female human subjects participated in the experiment. All had normal or corrected-to-normal vision. One subject was an experienced observer of visual psychophysics experiments, and three were naive to the purpose of the study. The study was conducted in accordance with the Declaration of Helsinki and approved by the Institutional Review Board at the University of Wisconsin-Madison. Written informed consent was obtained from all subjects before participation.

### Visual stimuli

Stimuli were presented on a 23-inch LED monitor (Planar SA2311W) operating at a refresh rate of 100 Hz and a spatial resolution of 1920 × 1080 pixels. The viewing distance was 60 cm. At this distance, 1° of visual angle corresponded to 39.5 pixels. Subjects viewed the display binocularly with head position stabilized by a chin rest and forehead support. Stimulus presentation and behavioral control were implemented in MATLAB (MathWorks, Natick, MA) using Psychophysics Toolbox 3 extensions (Brainard, 1997; Pelli, 1997). The monitor luminance output was linearized by gamma correction. The uniform display background was mid-gray, with a luminance of 40 cd/m². Eye position was monitored with a Tobii Pro Nano Eye Tracker (Tobii AB, Sweden) sampling at 60 Hz, with buffered gaze positions checked at 50 Hz (every two display frames).

The stimulus configuration is illustrated in Figure 1A. A stationary circular aperture, about 28° in diameter, contained a field of random dots. Eight smaller stationary circular apertures, each 5° in diameter, were positioned at 10° eccentricity and equally spaced around a central fixation point. Each smaller aperture contained a separate random-dot patch, and background dots were excluded from these regions. All dots were 4 pixels in diameter, had a luminance of 80 cd/m², and were presented at a density of 1.6 dots/deg². Round, anti-aliased dots were rendered using Psychtoolbox Screen(’DrawDots’) with alpha blending enabled. Dot positions were maintained at floating-point precision, allowing sub-pixel displacements and smoother motion rendering, particularly at low speeds. Each smaller patch was surrounded by a thin annular gap (0.5° in width) matching in luminance to the display background, spatially delineating all eight candidate regions. Because the same boundary surrounded every patch, target identity was defined solely by its speed difference from the background. During motion, dots reaching the boundary of an aperture wrapped around to the opposite side and continued moving in the same direction. The fixation point was 10 pixels in diameter and was presented at the center of the stationary circular aperture and the display.

**Figure 1.**
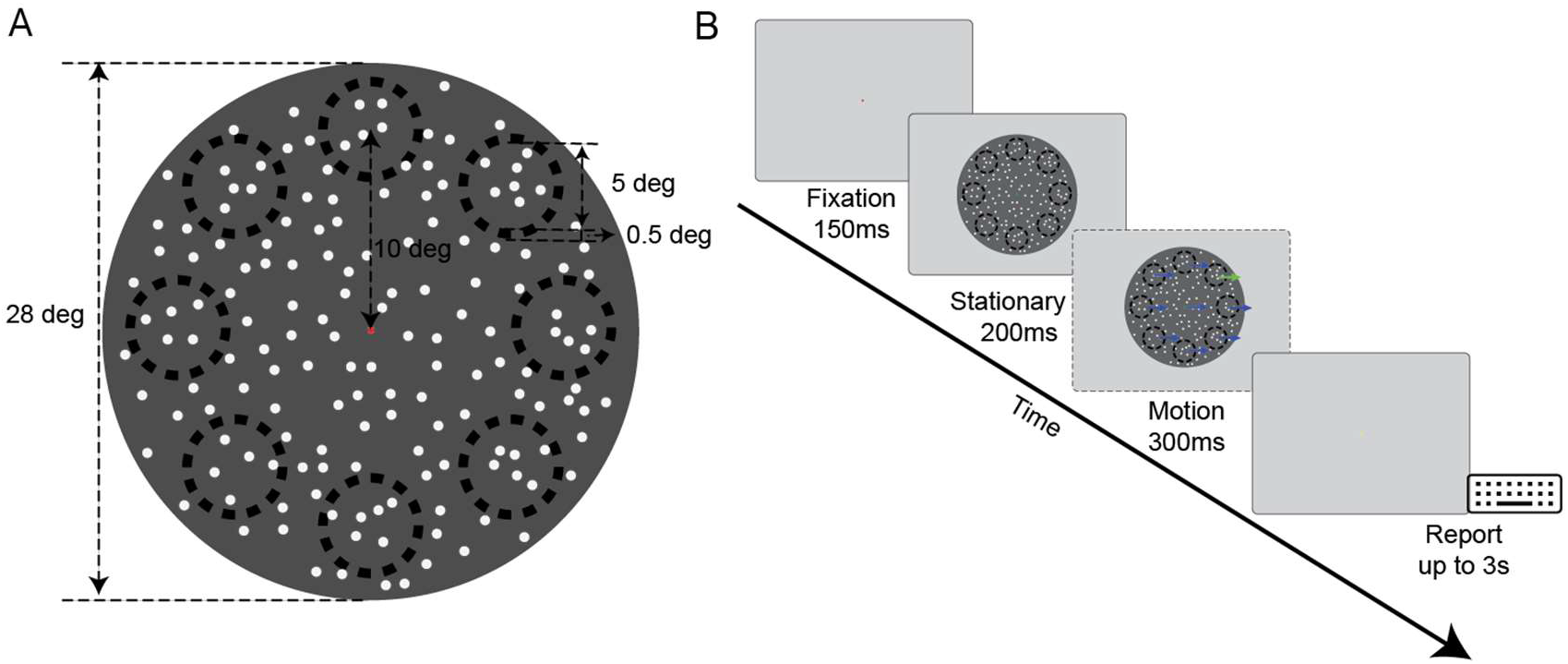
Stimulus configuration and trial sequence. (A) Spatial layout of the stimulus. A stationary circular aperture, 28° in diameter, contained the background random-dot pattern. Eight smaller circular apertures, each 5° in diameter, were positioned at 10° eccentricity and evenly spaced around a central fixation point. Each patch was surrounded by a thin annular gap (width 0.5°) matched in luminance to the display background. Background dots were excluded from the smaller apertures, each of which contained a separate random-dot patch. (B) Temporal sequence of a trial. Each trial began with a 150 ms of fixation, followed by 200 ms of stationary dots, 300 ms of motion, and a response interval of up to 3 s. The fixation point was red before fixation was acquired, green during the stationary and motion periods, and yellow during the response interval. On each trial, the background and all eight patches moved in the same direction. Seven patches moved at the background speed, whereas one target patch moved either faster (increment) or more slowly (decrement). The dark rings in the stimulus cartoon were for illustration purpose only.

### Experimental procedure

On each trial, the background and all eight patches moved in the same direction, either leftward or rightward. Seven patches moved at the same speed as the background, whereas one patch, the target, moved at a different speed. On increment trials, the target moved faster than the background; on decrement trials, it moved more slowly. The motion direction was either rightward (0°) or leftward (180°), and the target location among eight patches was selected randomly on each trial.

Two motion-coherence conditions were tested. In the 100% coherence condition, all dots moved coherently in the designated direction. In the reduced-coherence condition, 60% of the dots moved coherently, whereas the remaining 40% were replotted at random positions on each display frame. Signal and noisy dots assignments were randomized at each display frame. The same coherence level was applied to the background and all eight patches on a given trial.

The temporal sequence of each trial is illustrated in Figure 1B. Each trial began with a 150-ms fixation period, followed by 200 ms of stationary dots and 300 ms of motion. Subjects then had up to 3 s to respond. The fixation point indicated trial phase by color: red before fixation was acquired, green during the stationary and motion periods, and yellow during the response interval. Before each staircase run, the eye tracker was calibrated with a five-point procedure, with targets presented at the four corners and the center of the display (Tobii Pro SDK). Fixation was required throughout stimulus presentation (during the green phase). Binocular gaze position was monitored every other display frame (20ms). A trial was aborted right away if the gaze position of either eye deviated from the fixation point by more than 0.5°, or if no response was made within the response interval. Aborted trials were repeated with a newly randomized target location and motion direction.

Subjects performed an eight-alternative forced-choice (8AFC) localization task to detect which patch moved at a speed different from the background. To do so, subjects pressed one of eight spatially corresponding keys. When the target could not be identified confidently, subjects were instructed to make their best guess. No feedback was provided.

### Detection-threshold estimation

We used a staircase method to measure the minimum relative speed difference required to detect and localize the target. Detection thresholds were measured using a two-down, one-up adaptive staircase, which converges on approximately 70.7% correct performance (Levitt, 1971; Lu & Dosher, 2013). Target speed was adjusted relative to background speed in logarithmic steps, such that each step produced a fixed multiplicative change in speed. The initial step size was one-fifth of an octave, corresponding to a multiplicative factor of 2^1/5^ in speed. To refine sampling near threshold, the step size was halved after the first, third, and seventh reversals. Each staircase ended after 10 reversals, and the detection threshold was defined as the mean target speed over the final six reversals.

At the beginning of each increment staircase, target speed was selected randomly between two and four times the background speed. For decrement staircases, target speed was selected between one quarter and one half of the background speed. Thus, both staircase types began with a two- to four-fold relative-speed difference. These starting ranges were based on pilot testing, in which increment targets were readily detected over this range, whereas decrement performance was often poorer, particularly at low background speeds. The ranges were chosen to provide suprathreshold starting values while maintaining comparable relative-speed separations for increment and decrement staircases. Five background speeds were tested: 2.5, 5, 10, 20, and 30°/s. These speeds overlapped with those used in previous neurophysiological recordings from area MT (Huang et al., 2026), facilitating comparison between perceptual thresholds and the neural representation of multiple speeds.

Increment and decrement thresholds were measured with separate staircases at each background speed and coherence level. Increment thresholds were obtained at all five background speeds under both 100% and reduced coherence. Decrement thresholds were obtained at all five speeds under 100% coherence and at 5, 10, 20, and 30°/s under reduced coherence. The 2.5°/s decrement condition under reduced coherence could not be measured reliably because the target speed approached the lower limit of smooth apparent motion, and the staircase failed to converge in the presence of motion noise. This condition was therefore excluded from all analyses and figures.

Under 100% coherence, subjects completed at least 10 staircase runs for each combination of target condition and background speed. The final eight runs were analyzed, and preceding runs were treated as practice. Under reduced coherence condition, subjects completed at least six staircase runs for each condition, of which the final five were analyzed, and the preceding run was treated as practice. The detection threshold was computed as the arithmetic mean of the target speeds at the final six reversals. For each subject, all 100% coherence conditions were completed before the reduced-coherence conditions. Coherence was blocked, with each staircase run using a single coherence level throughout.

### Data analysis

Each staircase yielded the target speed required to detect and localize the relative-speed-defined target at the 70.7% correct criterion. Because speed is represented approximately logarithmically in area MT (Maunsell & Van Essen, 1983; Lisberger & Movshon, 1999; Nover et al., 2005), and perceptual sensitivity to speed is commonly expressed in relative rather than absolute units (Orban et al., 1985; De Bruyn & Orban, 1988), detection thresholds were quantified as speed ratios. The threshold ratio was defined generally as the faster speed divided by the slower speed. Thus, for the increment condition,

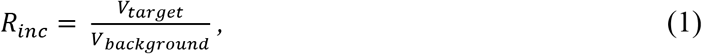

and for the decrement condition,

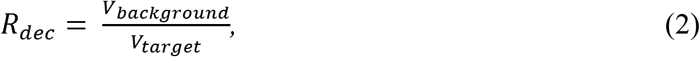

where *V_target_* and *V_background_* are the target and background speeds, respectively. Both ratios were therefore greater than 1. A larger ratio indicates that a greater relative-speed difference was required to detect and localize the target and thus indicates lower detection sensitivity.

Statistical analyses were performed on log-transformed threshold ratios, log_10_(*R*). We used the logarithmic transformation because it is appropriate for the multiplicative threshold measure, helps stabilize variance, and corresponds to the approximately logarithmic representation of speed in MT. Increment and decrement thresholds at different background speeds were obtained from separate staircase runs presented in randomized order. Consequently, threshold estimates were not paired across target conditions or background speeds. Analyses were performed separately for each subject, followed by group-level analyses across the four subjects.

To determine whether detection thresholds differed between increment and decrement targets and whether this difference depended on background speed, a two-way analysis of variance was conducted separately for each subject on log_10_(R), with target condition (increment versus decrement) and background speed as factors. Under 100% coherence, all five background speeds were included. Under reduced coherence, the analysis was restricted to the four speeds, 5, 10, 20, and 30°/s, at which thresholds were obtained for both conditions. Increment and decrement thresholds were also compared at each background speed using one-tailed Welch two-sample t-tests on log_10_(R). The one-tailed direction reflected the a priori prediction that decrement thresholds would exceed increment thresholds. The resulting (*p*) values were corrected across background speeds within each subject and coherence condition using the Benjamini-Hochberg false-discovery-rate procedure (Benjamini & Hochberg, 1995).

The magnitude of the increment-decrement difference was summarized using an asymmetry index (*ASI*) calculated from the mean threshold ratios for each subject, background speed, and coherence level:

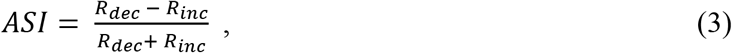

An index value of 0 indicates equal detection thresholds for increment and decrement targets, whereas a positive value indicates a higher threshold for decrement targets. Because increment and decrement thresholds were obtained from separate staircase runs, confidence intervals for the asymmetry index were estimated by independently resampling the increment and decrement runs with replacement. Bootstrap distributions were generated from 5,000 iterations, and 95% confidence intervals were defined by the 2.5th and 97.5th percentiles.

To quantify how the increment-decrement asymmetry changed with background speed, a linear model was fitted separately for each subject and coherence condition, with log_10_(R) as the dependent variable and target condition, log_10_-transformed background speed, and their interaction as predictors. The condition **×** log-speed interaction coefficient quantified the change in the increment-decrement threshold difference with increasing background speed. A negative coefficient indicates that the detection asymmetry decreased as background speed increased.

The effects of motion coherence were evaluated with a three-way analysis of variance conducted separately for each subject on log_10_(R), with target condition, coherence, and background speed as factors. This analysis was restricted to the balanced range of 5–30°/s, for which increment and decrement thresholds were available under both coherence conditions. As a more conservative confirmation, each subject was reduced to a single mean per condition and speed, and the group effects were tested across the four subjects with one-sample t tests.

The effect of reduced coherence was expressed as the ratio of the detection thresholds under 60% and 100% coherence:

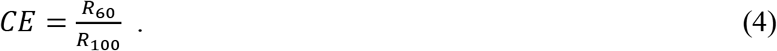

*R*_60_ and *R*_100_ were the mean threshold ratios under 60% and 100% coherence, respectively. A coherence effect (CE) value greater than 1 indicates higher thresholds under reduced coherence. Statistical effects of coherence were assessed using the three-way ANOVA on log_10_(R). On the logarithmic scale, the coherence contrast is log_10_(*CE*) = log_10_(*R_60_*) - log_10_(*R_100_*). The coherence main effect tested the overall effect of reduced coherence, while the coherence **×** speed and condition **×** coherence interactions tested whether this effect depended on background speed and target condition, respectively.

To characterize how relative-speed detection sensitivity scaled with background speed, Weber fractions (WF) were calculated with the background speed as the reference for both increment and decrement conditions:

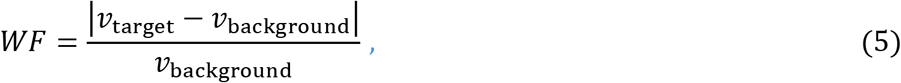

and log-transformed. Weber fractions were regressed on log-transformed background speed separately for each subject, with that subject’s staircase runs as independent observations. A slope of zero is consistent with Weber’s law, under which the Weber fraction is constant across speeds, whereas a negative slope indicates sub-Weber scaling, in which the Weber fraction decreases and relative sensitivity improves as background speed increases. The group slope was computed as the mean of the four individual-subject slopes, and its 95% confidence interval was obtained across the four subjects using a t distribution (df = 3). The full coherence increment condition was treated separately because its Weber fraction followed the non-monotonic, U-shaped function of speed described in earlier work (De Bruyn & Orban, 1988), decreasing to a minimum near 5°/s before rising to a plateau. The regression was therefore restricted to the 10–30°/s plateau to test for Weber’s Law, and the minimum at 5°/s was tested separately with one-tailed Welch t tests comparing the Weber fraction at 5°/s with those at 2.5 and 10°/s, corrected by the false discovery rate. The one-tailed direction followed the previously reported location of this minimum.

All analyses were performed in MATLAB (MathWorks). The significance threshold was α = 0.05.

## Results

We tested whether targets moving faster and slower than a moving background differed in detectability. Four subjects performed an 8AFC localization task, and separate adaptive staircases estimated increment and decrement thresholds across five background speeds under 100% and reduced motion coherence.

### Detection thresholds were higher for slower than faster targets

Detection thresholds were expressed as speed ratios, with larger ratios indicating that a greater relative-speed difference was required to detect and localize the target. Figure 2 shows the threshold ratios for each subject as a function of background speed under 100% (Fig. 2A1-D1) and reduced coherence (Fig. 2A2-D2). Across all four subjects, thresholds were higher for decrement targets than for increment targets. Thus, a target moving more slowly than the background required a larger relative-speed difference for detection than a target moving faster. Under 100% coherence, the main effect of target condition was significant for every subject (two-way analysis of variance on log-transformed ratios, F(1,70) = 38.7-152.2, all p < 0.001, partial η² = 0.36-0.69). The condition effect was also significant under reduced coherence (F(1,32) = 111.4-274.0, all p < 0.001, partial η² = 0.78-0.90) (Table 1).

**Figure 2.**
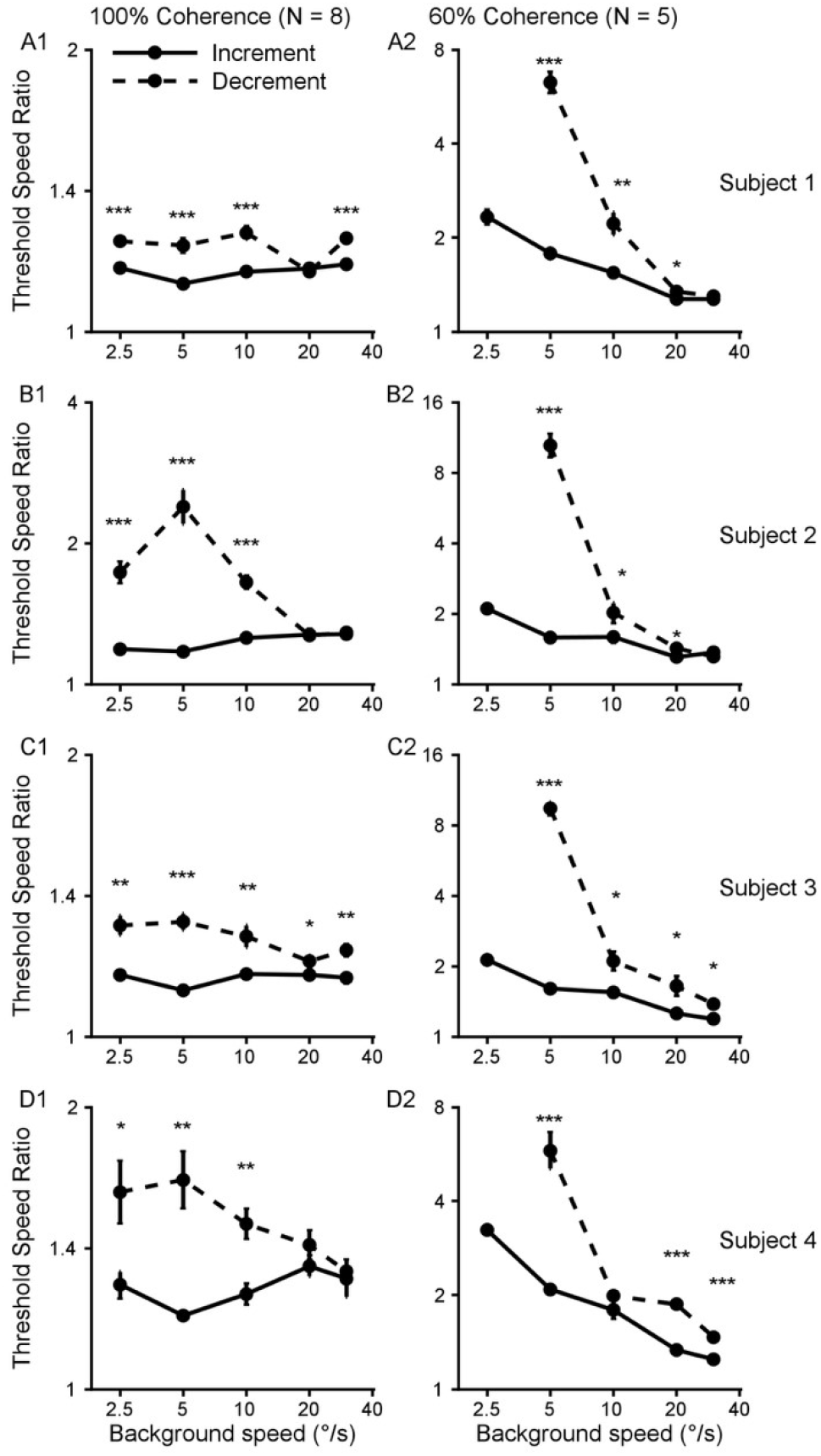
Relative-speed detection thresholds for individual subjects. Threshold speed ratios (either 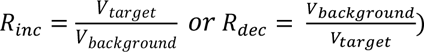 are plotted as a function of background speed for each of the four subjects (rows) under 100% coherence (A1–D1; eight staircase runs per condition) and 60% coherence (A2–D2; five runs per condition). Both axes are logarithmic. Solid lines show increment thresholds, and dashed lines show decrement thresholds. Symbols indicate mean threshold ratios across runs; error bars show SEM. Larger ratios indicate that a greater relative-speed difference was required to detect and localize the target. Asterisks indicate significant differences between increment and decrement thresholds at individual background speeds (one-tailed Welch two-sample t tests on log-transformed ratios, with false-discovery-rate correction across speeds within each subject and coherence condition; *p<0.05, **p<0.01, ***p<0.001). The decrement threshold at 2.5°/s under 60% coherence could not be measured reliably and is omitted. Subject-specific mean threshold ratios, SEMs, and statistical comparisons are reported in Supplementary Table 1.

**Table 1.** Effects of target condition and its interaction with background speed on relative-speed detection thresholds. Two-way ANOVAs were conducted separately for each subject on log-transformed threshold ratios under 100% and reduced coherence, with target condition (increment versus decrement) and background speed as factors. The table shows the main effect of target condition and the target-condition **×** background-speed interaction. Partial η²ₚ indicates effect size.

|  |  |  | **Subject 1** |  | **Subject 2** |  | **Subject 3** |  | **Subject 4** |  |
|---|---|---|---|---|---|---|---|---|---|---|
| **Effects** | **df** | ***p*** | **F** | **η<sup>2</sup><sub>p</sub>** | **F** | **η<sup>2</sup><sub>p</sub>** | **F** | **η<sup>2</sup><sub>p</sub>** | **F** | **η<sup>2</sup><sub>p</sub>** |
| **100% coherence** |  |  |  |  |  |  |  |  |  |  |
| **Condition** | 1, 70 | <0.001 *** | 89.0 | 0.56 | 152.2 | 0.69 | 94.8 | 0.58 | 38.7 | 0.36 |
| **Condition × Speed** | 4, 70 | <0.01 † ** | 8.1 | 0.32 | 35.6 | 0.67 | 5.3 | 0.23 | 5.0 | 0.22 |
| **60% coherence** |  |  |  |  |  |  |  |  |  |  |
| **Condition** | 1, 32 | <0.001 *** | 180.0 | 0.85 | 175.1 | 0.85 | 274.0 | 0.90 | 111.4 | 0.78 |
| **Condition × Speed** | 3, 32 | <0.001 *** | 83.3 | 0.89 | 121.9 | 0.92 | 104.2 | 0.91 | 30.0 | 0.74 |

† 100% Condition × Speed: Subjects 1–3 (p < 0.001), Subject 4 (p = 0.0013).

Comparisons at individual background speeds supported the same conclusion. Under 100% coherence, decrement thresholds significantly exceeded increment thresholds at most low and intermediate background speeds (Fig. 2, Welch t-tests, false discovery rate corrected). The difference generally diminished at higher background speeds, becoming nonsignificant in three of the four subjects at 20°/s and in two of the four subjects at 30°/s. Under reduced coherence, decrement thresholds were substantially higher at the lowest measured background speeds, with the difference again becoming smaller as background speed increased. Threshold ratios and statistical comparisons for each subject and background speed are reported in Supplementary Table 1.

These results demonstrate a robust asymmetry in relative-speed detection: slower-than-background targets were generally more difficult to detect than faster-than-background targets.

### The detection asymmetry decreased with background speed

The difference between increment and decrement thresholds depended systematically on background speed. The target-condition **×** speed interaction was significant for all subjects under both 100% coherence (F(4,70) = 5.0-35.6, all p < 0.001), and reduced coherence (F(3,32) = 30.0-121.9, all p < 0.001) (Table 1).

This dependence is summarized by ASI in Figure 3. An ASI of 0 indicates equal increment and decrement thresholds, whereas positive values indicate higher thresholds for decrement targets. For every subject, the ASI was larger at the lower background speeds and generally approached 0 as background speed increased (Fig. 3A, B). Thus, the perceptual disadvantage for slower targets was most pronounced at low speeds and diminished at higher speeds.

**Figure 3.**
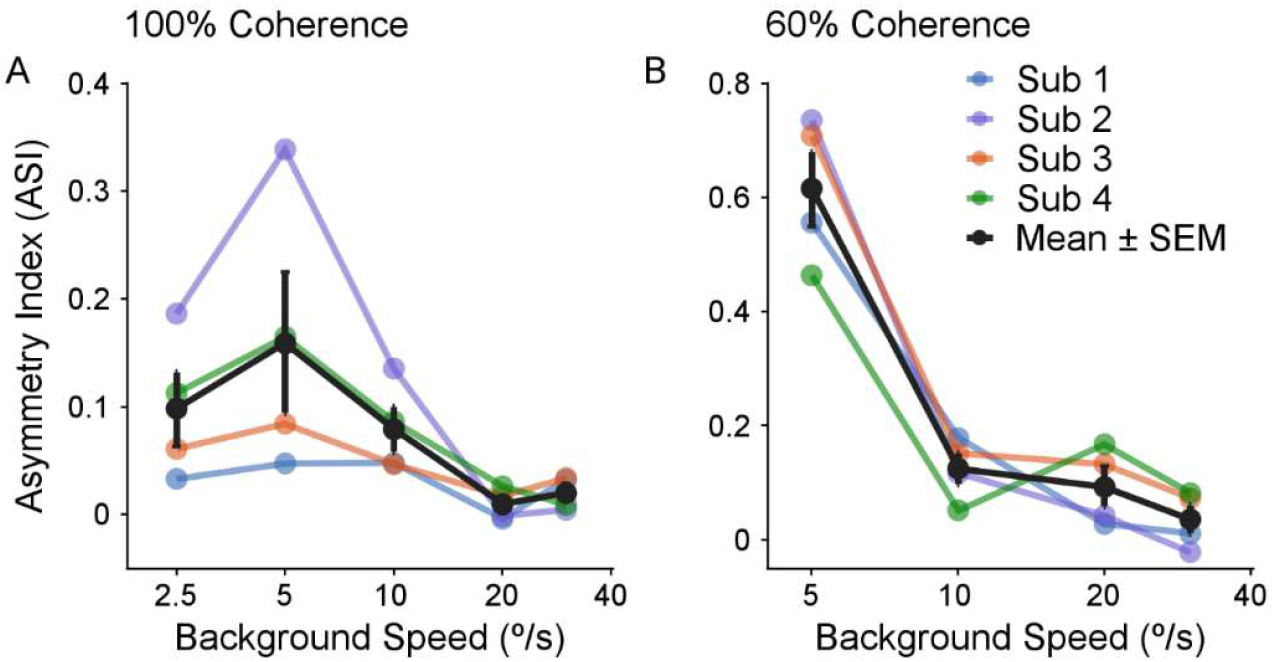
Asymmetry between increment and decrement detection thresholds. The asymmetry index (ASI) is plotted as a function of background speed under 100% coherence (A) and 60% coherence (B). ASI index was calculated from the mean increment and decrement threshold ratios across staircase runs at each background speed. A value of 0 indicates equal thresholds, whereas positive values indicate higher thresholds for decrement than increment targets. Colored lines show individual subjects; the black line shows the mean across four subjects, with error bars indicating SEM. No ASI is shown at 2.5°/s under 60% coherence because the decrement threshold could not be measured reliably. Bootstrap 95% confidence intervals for individual-subject ASIs are reported in Supplementary Table 2.

The decline in asymmetry was steeper under reduced coherence (Fig. 3B). To quantify the speed dependence within each coherence condition, we fitted a linear model to the log-transformed threshold ratios for each subject. The target-condition × log_10_(background-speed) interaction coefficient was negative for all subjects under reduced coherence, ranging from −0.40 to −1.02 (Supplementary Table 3), and was significant across subjects (p = 0.011) (Supplementary Table 4). Under 100% coherence, the coefficients were smaller, ranging from −0.02 to −0.23; although three subjects individually showed significant negative coefficients, the effect was not significant across subjects (p = 0.13) (Supplementary Tables 3 and 4).

The increment-decrement asymmetry therefore depended on both motion coherence and background speed. It was strongest when motion coherence and background speed were low and became progressively smaller as background speed increased.

### Reduced coherence preferentially impaired detection of slower targets

Reducing motion coherence from 100% to 60% increased detection thresholds, but the effect was not uniform across target conditions or background speeds. Figure 4 shows the coherence effect for increment and decrement targets. Reduced coherence generally elevated thresholds for both conditions, but the increase was consistently larger for decrement targets and at lower background speeds.

**Figure 4.**
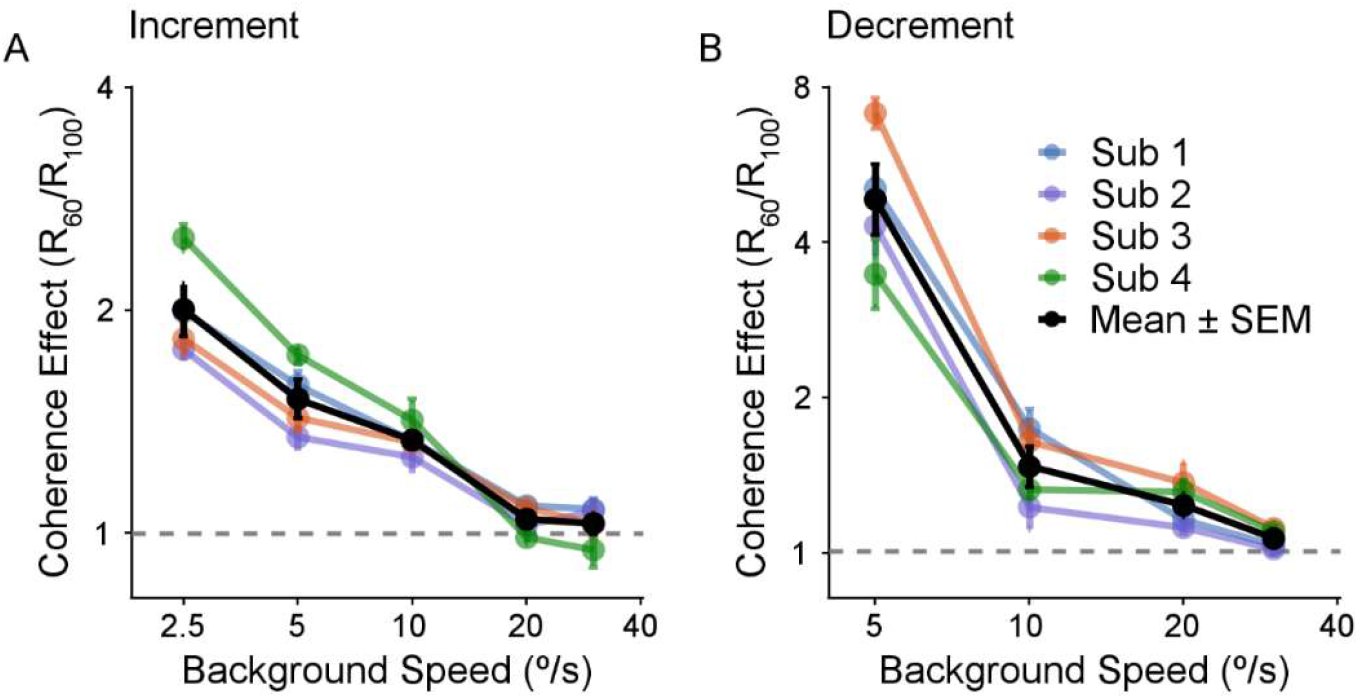
Effect of reduced motion coherence on relative-speed detection thresholds. The coherence effect, *R*_60_/*R*_100_, is plotted as a function of background speed for increment (A) and decrement (B) targets. Both axes are logarithmic. A value of 1 indicates no effect of coherence, whereas values greater than 1 indicate that reduced coherence increased the relative-speed difference required for target detection. Colored lines show individual subjects; the black line shows the mean across four subjects. For individual subjects, error bars reflect the propagated SEMs from the threshold ratios under the two coherence conditions; for the group mean, error bars indicate SEM across subjects. The decrement condition begins at 5°/s background speed because the threshold at 2.5°/s under 60% coherence could not be measured reliably.

A three-way analysis of variance was conducted separately for each subject over the balanced range of 5-30°/s, with target condition, coherence, and background speed as factors. The main effect of coherence was significant in every subject (F(1,88) = 185.7-958.4, all p < 0.001, partial η² = 0.68-0.92), confirming that reduced coherence increased the relative-speed difference required for target detection.

The target condition **×** coherence interaction was also significant for all subjects (F(1,88)=36.1-276.4, all p < 0.001, partial η² = 0.29-0.76). Thus, reducing coherence impaired detection of slower targets more strongly than detection of faster targets. The effect of reduced coherence was greatest at low background speeds and diminished as speed increased; the coherence **×** speed interaction was significant as well (F(3,88) = 78.6-294.5, all p < 0.001, partial η² = 0.73-0.91).

Finally, the target condition **×** coherence **×** speed interaction was significant in every subject (F(3,88) = 13.3-125.4, all p < 0.001), partial η² = 0.31-0.81). The disproportionate impairment of decrement detection by reduced coherence was therefore strongest at low background speeds. Conservative analyses across the four subjects confirmed the overall coherence effect (p = 0.002), the target condition **×** coherence interaction (p = 0.009), coherence **×** speed interaction (p < 0.001), and the three-way condition **×** coherence **×** speed interaction (p = 0.018) (Supplementary Table 4).

Together, these results show that motion noise did not merely produce a uniform loss of sensitivity. It preferentially impaired detection of slower-than-background targets, particularly at low speeds, thereby amplifying the asymmetry between increment and decrement detection.

### Weber scaling of relative-speed detection across background speeds

To characterize how sensitivity scaled with background speed, Weber fractions were plotted as a function of background speed and analyzed on logarithmic axes (Fig. 5; Individual-subject regression slopes and 95% confidence intervals are reported in Supplementary Table 5). A slope of zero indicates a constant Weber fraction and is therefore consistent with Weber’s law, whereas a negative slope indicates sub-Weber scaling, in which relative sensitivity improves with increasing background speed.

**Figure 5.**
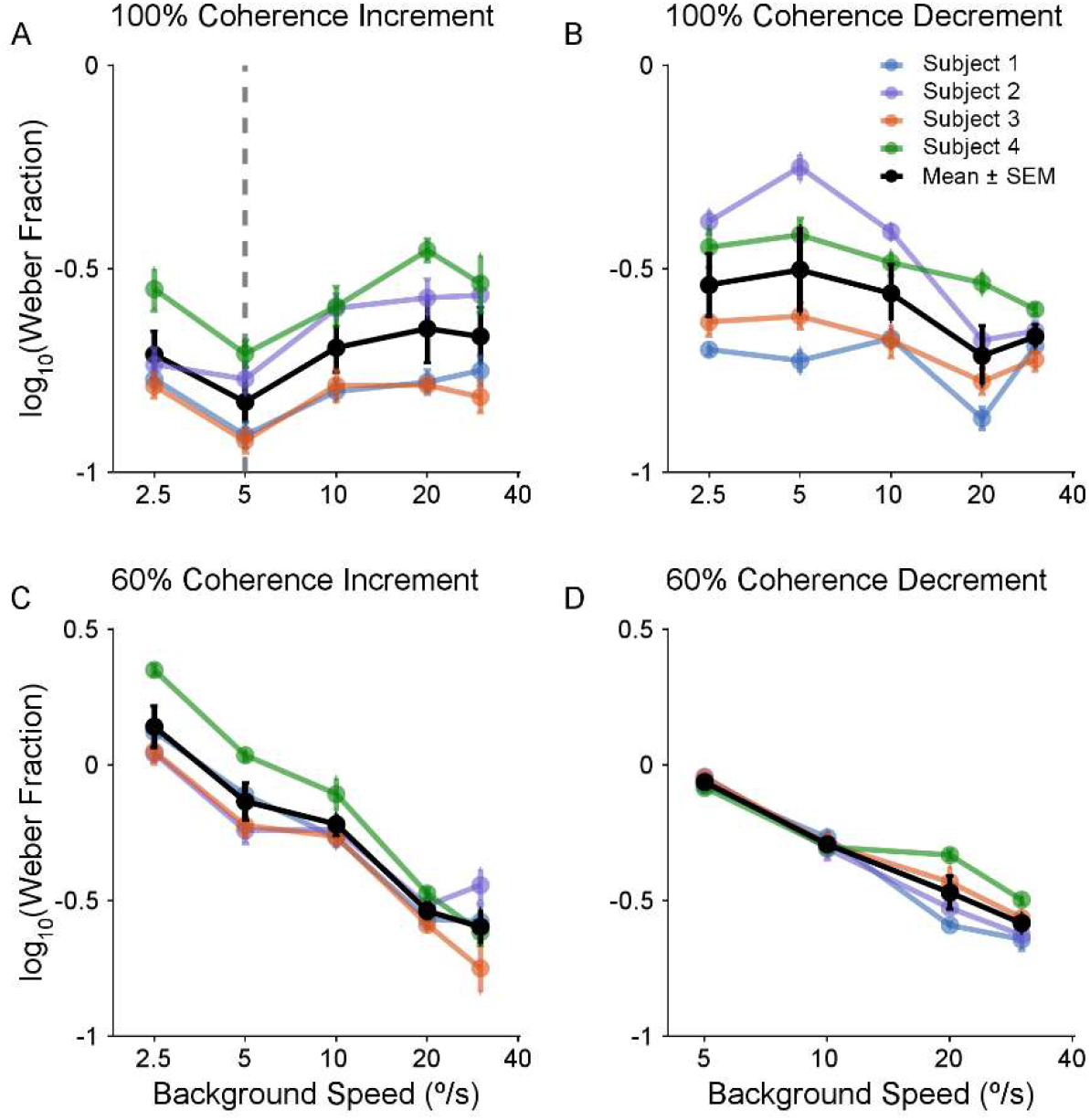
Weber scaling of relative-speed detection across background speed. The log_10_ of the Weber fraction is plotted as a function of background speed on a logarithmic x-axis for four conditions: 100% coherence increment (A), 100% coherence decrement (B), 60% coherence increment (C), and 60% coherence decrement (D). Weber fraction is defined as 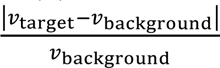. Colored lines show individual subjects, with error bars indicating SEM across staircase runs. The black line shows the mean across subjects, with error bars indicating SEM of the mean across the four subjects. In A, the dashed vertical line marks 5°/s, where the 100%-coherence increment Weber fraction reached a local minimum. The 2.5°/s decrement condition under 60% coherence is absent from D because it could not be measured reliably. Individual-subject regression slopes and 95% confidence intervals are reported in Supplementary Table 5.

Under 100% coherence, increment detection showed a non-monotonic dependence on background speed (Fig. 5A). The Weber fraction reached a minimum near 5°/s and was relatively constant from 10 to 30°/s. Over this 10-30°/s range, the group regression slope did not differ from zero (0.07, 95% CI [−0.07, 0.21]), consistent with Weber’s law. The Weber fraction at 5°/s was significantly lower than at 10°/s in all four subjects and lower than at 2.5°/s in three of the four subjects (Fig. 5A, vertical dashed line).

Under 100% coherence, decrement detection showed no clear monotonic scaling with background speed (Fig. 5B). Although the group regression slope was negative, its confidence interval included zero (−0.17, 95% CI [−0.37, 0.03]), indicating no reliable overall change in Weber fraction with increasing background speed.

In contrast, under reduced coherence, Weber fractions decreased systematically with increasing background speed for both increment and decrement detection (Fig. 5C, D). Regression slopes were negative in every subject, and the group confidence intervals excluded zero for both increment (group slope −0.68, 95% CI [−0.96, −0.41]) and decrement (−0.66, 95% CI [−0.88, −0.44]) conditions. Thus, both conditions showed clear sub-Weber scaling under reduced coherence, with similar estimated group slopes.

## Discussion

We found a consistent asymmetry in relative-speed detection: targets moving more slowly than the background required larger relative-speed differences for detection than targets moving faster. The asymmetry was found in all four subjects, was largest at low background speeds, and declined as speed increased. Reducing motion coherence from 100% to 60% raised thresholds in both increment and decrement conditions, but the increase was greater for decrement targets, especially at low speeds, so the asymmetry became stronger under motion noise. Under 100% coherence, increment detection was approximately Weber-like from 10 to 30°/s, whereas under reduced coherence both increment and decrement Weber fractions decreased with increasing background speed. Relative-speed sensitivity therefore depended on the sign of the target-background speed difference, background speed, and motion coherence.

### Faster-target advantage in relative-speed detection

Our results agree with Sekuler (1990) in showing a faster-target advantage at low background speeds. The comparison also reveals that the apparent pattern of asymmetry depends on how relative-speed thresholds are expressed. With background-referenced Weber fractions, our 100%-coherence data showed evidence of a high-speed reversal, with decrement Weber fractions falling below increment Weber fractions in several subjects (Fig. 5A, B). With the symmetric faster/slower speed ratio used for our primary analyses, however, the asymmetry declined toward zero at high speeds without a consistent reversal (Fig. 3A). Thus, the qualitative appearance of a crossover depends in part on the threshold metric, even though both measures show a strong faster-target advantage at low speeds.

The effect of motion noise also parallels Sekuler’s findings. In both studies, making the motion signal less reliable increased the increment-decrement asymmetry. Our results extend this effect across a broader, systematically sampled range of background speeds and in a near peripheral eight-location localization task. Unlike Sekuler, we also observed a clear asymmetry under full coherence, indicating that added motion noise is not required for the faster-target advantage, although it substantially amplifies it.

Related findings further indicate that there is no universal advantage for either speed increments or decrements. Ivry and Cohen (1992) reported a faster-target advantage in visual search, whereas Hohnsbein and Mateeff (2002) found greater sensitivity to temporal speed decrements under conditions in which aperture geometry and visible motion path were important. Together, these findings suggest that speed asymmetries depend on task, stimulus configuration, speed range, and motion reliability.

### Relation to speed weighting in MT and natural-scene statistics

The behavioral pattern is consistent with a biased faster-speed weighting previously observed in area MT. When two overlapping speed components were presented simultaneously within the RFs of MT neurons, neural responses were weighted more toward the faster component, and the bias was strongest at relatively low speeds (Huang et al., 2026). In a related study, preliminary findings reported in a conference abstract suggest that a similar faster-speed bias at low speeds (<20°/s) also occurred for spatially separated stimuli (Chakrala et al., 2023). If relative-speed detection depends on this neural representation, a faster target should be represented more strongly relative to its background than an equally separated slower target. The correspondence extends beyond the direction of the bias to its speed dependence: both the perceptual asymmetry and the neural faster-speed bias were strongest at low speeds. Our psychophysical data do not establish that MT faster-speed weighting causes the perceptual asymmetry, but they are consistent with that mechanism. Sekuler’s (1990) sigmoidal speed-response model provides one alternative explanation, while Ivry and Cohen (1992) suggested that high-pass filtering in the velocity domain might contribute to faster-target asymmetries in visual search. The present results do not uniquely identify the mechanism, but they raise the possibility that biased weighting of multiple speed signals within MT receptive fields contributes to the observed perceptual asymmetry.

Natural-scene statistics provide a possible functional context. Figural regions in dynamic natural scenes tend to move faster than their backgrounds (Friedman et al., 2025). Greater weighting of faster motion would therefore emphasize a signal statistically associated with figure-ground assignment. In our task, faster-than-background targets were detectable at smaller relative-speed differences than slower-than-background targets, as expected if the visual system exploits this natural-scene statistical regularity. This correspondence suggests a possible role for faster-speed weighting in motion-based figure-ground segregation.

### Motion noise amplifies the asymmetry

Reducing coherence raised thresholds for both target types, but it did not produce a uniform reduction of sensitivity. The increase was substantially larger for decrement targets and at lower background speeds, and the target-condition **×** coherence and three-way interactions were significant in every subject (Table 2). Motion noise therefore made the faster-target advantage stronger rather than weaker. The main effect of target condition accounted for a larger share of the variance under reduced coherence (partial η² = 0.78-0.90) than under 100% coherence (0.36-0.69; Table 1), even though the reduced-coherence analysis excluded the lowest background speed, where asymmetries tended to be large. The decline of the asymmetry with background speed was also steeper under reduced coherence (Fig. 3B).

**Table 2.** Effects involving motion coherence on relative-speed detection thresholds. Three-way ANOVAs were conducted separately for each subject on log-transformed threshold ratios, with target condition, motion coherence, and background speed as factors. Analyses were restricted to the balanced range of 5–30°/s, for which both increment and decrement thresholds were available under both 100% and 60% coherence conditions. Only effects involving coherence are shown; the effects of target condition and its interaction with background speed are reported in Table 1. Partial η²ₚ indicates effect size.

|  | **for all subjects** | **for all subjects** | **Subject 1** |  | **Subject 2** |  | **Subject 3** |  | **Subject 4** |  |
|---|---|---|---|---|---|---|---|---|---|---|
| **Effects** | **df** | ***p*** | **F** | **η<sup>2</sup><sub>p</sub>** | **F** | **η<sup>2</sup><sub>p</sub>** | **F** | **η<sup>2</sup><sub>p</sub>** | **F** | **η<sup>2</sup><sub>p</sub>** |
| **Coherence** | 1, 88 | <0.001 *** | 958.4 | 0.92 | 185.7 | 0.68 | 800.0 | 0.90 | 243.2 | 0.73 |
| **Condition × Coherence** | 1, 88 | <0.001 *** | 190.0 | 0.68 | 43.6 | 0.33 | 276.4 | 0.76 | 36.1 | 0.29 |
| **Coherence × Speed** | 3, 88 | <0.001 *** | 294.5 | 0.91 | 78.6 | 0.73 | 232.8 | 0.89 | 79.8 | 0.73 |
| **Condition × Coherence × Speed** | 3, 88 | <0.001 *** | 109.0 | 0.79 | 43.8 | 0.60 | 125.4 | 0.81 | 13.3 | 0.31 |

Because target and background were degraded by the same coherence manipulation, a uniform loss of motion information would not by itself predict the selective impairment of slower targets. Coherence-dependent changes in perceived speed or speed uncertainty may contribute to performance (Chuang et al., 2016; Weiss et al., 2002; Stocker & Simoncelli, 2006; Zhang & Stocker, 2022), but an effect common to target and background likewise would not be expected to favor faster targets. Instead, the speed difference that distinguishes a slower target from its background appears to be more fragile than the one that distinguishes a faster target. Noise exposed and enlarged a bias already present under 100% coherence.

This amplification may matter for figure-ground segregation in natural vision, where motion signals are often noisy, sparse, or partially occluded. If segregation of moving regions reflects the regularity that figures tend to move faster than their backgrounds (Friedman et al., 2025), the influence of that regularity would be expected to grow as sensory evidence weakens, in line with the general finding that prior expectations exert a greater influence on speed perception when stimulus reliability is low (Weiss et al., 2002; Stocker & Simoncelli, 2006). Faster-than-background regions would then remain readily segregated under noisy conditions, whereas slower-than-background regions would require substantially larger speed differences to be detected. Measuring thresholds across a wider range of coherence levels would show whether the asymmetry grows continuously as motion signals weaken.

### Scaling of relative-speed sensitivity with background speed

Under 100% coherence, increment detection showed its best relative sensitivity near 5°/s and was approximately constant from 10 to 30°/s, consistent with Weber’s law over that range. This resembles classic speed-discrimination results, in which sensitivity is best at intermediate speeds and becomes approximately Weber-like over part of the tested range (Orban et al., 1985; De Bruyn & Orban, 1988). Decrement detection under 100% coherence, however, showed no reliable monotonic scaling with speed.

Under reduced coherence, Weber fractions decreased with increasing background speed for both increments and decrements, producing negative slopes of similar magnitude. These slopes parallel the coherence **×** speed interaction: because reduced coherence elevated thresholds most at low background speeds, relative sensitivity under noise improved as background speed increased. Relative-speed detection therefore did not follow a single Weber scaling across conditions. This is consistent with evidence that speed-based segmentation and conventional speed discrimination need not have the same speed dependence (Masson et al., 1999). Here, scaling within a single segmentation task also changed with motion coherence and with whether the target was faster or slower than the background.

### Limitations and conclusion

Several limitations should be considered in interpreting these findings. Only four observers were tested, although the main effects were consistent across subjects. The candidate regions were spatially delineated by identical annular gaps, so relative speed defined target identity rather than the figure boundary itself. The proposed link to MT is also based on correlation with previous neurophysiological findings rather than causal manipulation. Within these limits, the behavioral pattern is consistent: faster-than-background targets are detected at smaller relative-speed differences than slower-than-background targets, with the largest asymmetry at low speeds and under reduced motion coherence. Its correspondence with faster-speed weighting in MT and with the natural tendency for figural regions to move faster than their backgrounds suggests that relative-speed sensitivity may contribute to selecting motion signals useful for figure–ground segregation.

## Acknowledgments

We thank Bikalpa Ghimire and Steven Wiesner for helpful discussions and assistance during this project, Yuning Jiang for assistance with the data collection, and Abhi Padala for helping with setting up the psychophysics rig and computer programming.

## Use of generative AI and AI-assisted technologies

During the preparation of this manuscript, the authors used Claude (Anthropic) and ChatGPT (OpenAI) to assist with language editing and to improve the clarity, readability, and organization of the text. Claude was also used to assist with generating code for calculating Weber fractions and conservative group-level analyses. The authors reviewed and revised all AI-assisted text and validated and modified the generated code to ensure accuracy. The authors take full responsibility for the content, accuracy, and integrity of the work.

---

**STable 1.** Relative-speed detection threshold ratios for each subject, target condition, background speed, and motion-coherence level. Values are means ± SEM across staircase runs (N = 8 under 100% coherence; N = 5 under 60% coherence) and correspond to the data shown in Figure 2. Increment and decrement thresholds were compared at each background speed using one-tailed Welch two-sample t-tests on log10-transformed threshold ratios, with Benjamini–Hochberg false-discovery-rate correction across background speeds within each subject and coherence condition. The 2.5°/s decrement threshold under 60% coherence could not be measured reliably and is therefore not reported. *p < 0.05, **p < 0.01, ***p < 0.001.

|  |  | **100% coherence** (N = 8 staircase runs) |  |  | **60% coherence** (N = 5 staircase runs) |  |  |
|---|---|---|---|---|---|---|---|
|  | **Speed**<br>(°/s) | **Increment**<br>(mean±SEM) | **Decrement**<br>(mean±SEM) | ***p*** | **Increment**<br>(mean± SEM) | **Decrement**<br>(mean±SEM) | ***p*** |
| **Subject 1** | 2.5 | 1.171 ± 0.009 | 1.251 ± 0.008 | <0.001<br>*** | 2.341 ± 0.122 | — | — |
|  | 5 | 1.125 ± 0.008 | 1.238 ± 0.020 | <0.001<br>*** | 1.787 ± 0.072 | 6.354 ± 0.494 | <0.001<br>*** |
|  | 10 | 1.160 ± 0.010 | 1.277 ± 0.019 | <0.001<br>*** | 1.548 ± 0.048 | 2.250 ± 0.198 | 0.009<br>** |
|  | 20 | 1.169 ± 0.011 | 1.159 ± 0.010 | 0.73 | 1.273 ± 0.019 | 1.344 ± 0.010 | 0.011<br>* |
|  | 30 | 1.181 ± 0.012 | 1.259 ± 0.012 | <0.001<br>*** | 1.273 ± 0.036 | 1.300 ± 0.029 | 0.28 |
| **Subject 2** | 2.5 | 1.189 ± 0.018 | 1.749 ± 0.089 | <0.001<br>*** | 2.106 ± 0.065 | — | — |
|  | 5 | 1.174 ± 0.014 | 2.449 ± 0.191 | <0.001<br>*** | 1.586 ± 0.066 | 10.701 ± 1.114 | <0.001<br>*** |
|  | 10 | 1.257 ± 0.018 | 1.654 ± 0.045 | <0.001<br>*** | 1.598 ± 0.078 | 2.061 ± 0.210 | 0.044<br>* |
|  | 20 | 1.279 ± 0.033 | 1.273 ± 0.022 | 0.54 | 1.309 ± 0.031 | 1.423 ± 0.012 | 0.019<br>* |
|  | 30 | 1.280 ± 0.026 | 1.290 ± 0.016 | 0.45 | 1.374 ± 0.053 | 1.312 ± 0.022 | 0.83 |
| **Subject 3** | 2.5 | 1.165 ± 0.010 | 1.318 ± 0.033 | 0.002<br>** | 2.139 ± 0.114 | — | — |
|  | 5 | 1.121 ± 0.008 | 1.329 ± 0.025 | <0.001<br>*** | 1.607 ± 0.060 | 9.509 ± 0.591 | <0.001<br>*** |
|  | 10 | 1.167 ± 0.013 | 1.284 ± 0.037 | 0.007<br>** | 1.549 ± 0.037 | 2.135 ± 0.183 | 0.013<br>* |
|  | 20 | 1.164 ± 0.005 | 1.205 ± 0.017 | 0.021<br>* | 1.258 ± 0.014 | 1.673 ± 0.152 | 0.019<br>* |
|  | 30 | 1.157 ± 0.014 | 1.238 ± 0.016 | 0.002<br>** | 1.195 ± 0.046 | 1.379 ± 0.031 | 0.013<br>* |
| **Subject 4** | 2.5 | 1.298 ± 0.045 | 1.658 ± 0.135 | 0.017<br>* | 3.239 ± 0.080 | — | — |
|  | 5 | 1.199 ± 0.015 | 1.703 ± 0.125 | 0.002<br>** | 2.087 ± 0.044 | 6.007 ± 0.783 | <0.001<br>*** |
|  | 10 | 1.266 ± 0.030 | 1.509 ± 0.055 | 0.002<br>** | 1.807 ± 0.104 | 1.996 ± 0.058 | 0.082 |
|  | 20 | 1.355 ± 0.021 | 1.432 ± 0.053 | 0.12 | 1.336 ± 0.011 | 1.876 ± 0.056 | <0.001<br>*** |
|  | 30 | 1.323 ± 0.066 | 1.336 ± 0.012 | 0.36 | 1.247 ± 0.027 | 1.468 ± 0.020 | <0.001<br>*** |

**STable 2.** Asymmetry index (ASI) for each subject and background speed under 100% and 60% motion coherence. ASI was calculated from the mean increment and decrement threshold ratios as *ASI* = 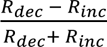. Values are point estimates with bootstrap 95% confidence intervals in brackets, obtained by independently resampling increment and decrement staircase runs with replacement (5,000 iterations). Positive values indicate higher decrement than increment thresholds; negative values indicate the reverse. The 2.5°/s decrement threshold under 60% coherence could not be measured reliably, so no ASI is reported for that condition.

|  | **Speed (°/s)** | **100% coherence** | **60% coherence** |
|---|---|---|---|
| **Subject 1** | 2.5 | 0.033 [0.024, 0.042] | — |
|  | 5 | 0.048 [0.032, 0.065] | 0.561 [0.512, 0.614] |
|  | 10 | 0.048 [0.032, 0.063] | 0.185 [0.110, 0.262] |
|  | 20 | -0.004 [-0.015, 0.008] | 0.027 [0.014, 0.042] |
|  | 30 | 0.032 [0.019, 0.046] | 0.011 [-0.022, 0.041] |
| **Subject 2** | 2.5 | 0.191 [0.145, 0.238] | — |
|  | 5 | 0.352 [0.284, 0.407] | 0.742 [0.689, 0.781] |
|  | 10 | 0.136 [0.108, 0.163] | 0.126 [0.038, 0.224] |
|  | 20 | -0.002 [-0.033, 0.024] | 0.042 [0.020, 0.064] |
|  | 30 | 0.004 [-0.020, 0.025] | -0.023 [-0.059, 0.012] |
| **Subject 3** | 2.5 | 0.062 [0.039, 0.087] | — |
|  | 5 | 0.085 [0.065, 0.101] | 0.711 [0.678, 0.740] |
|  | 10 | 0.048 [0.022, 0.077] | 0.159 [0.081, 0.232] |
|  | 20 | 0.017 [0.005, 0.032] | 0.142 [0.060, 0.215] |
|  | 30 | 0.034 [0.017, 0.049] | 0.072 [0.031, 0.108] |
| **Subject 4** | 2.5 | 0.122 [0.043, 0.200] | — |
|  | 5 | 0.174 [0.109, 0.242] | 0.484 [0.384, 0.562] |
|  | 10 | 0.088 [0.048, 0.128] | 0.050 [-0.001, 0.109] |
|  | 20 | 0.028 [-0.003, 0.069] | 0.168 [0.146, 0.197] |
|  | 30 | 0.005 [-0.044, 0.044] | 0.081 [0.058, 0.103] |

**STable 3.**
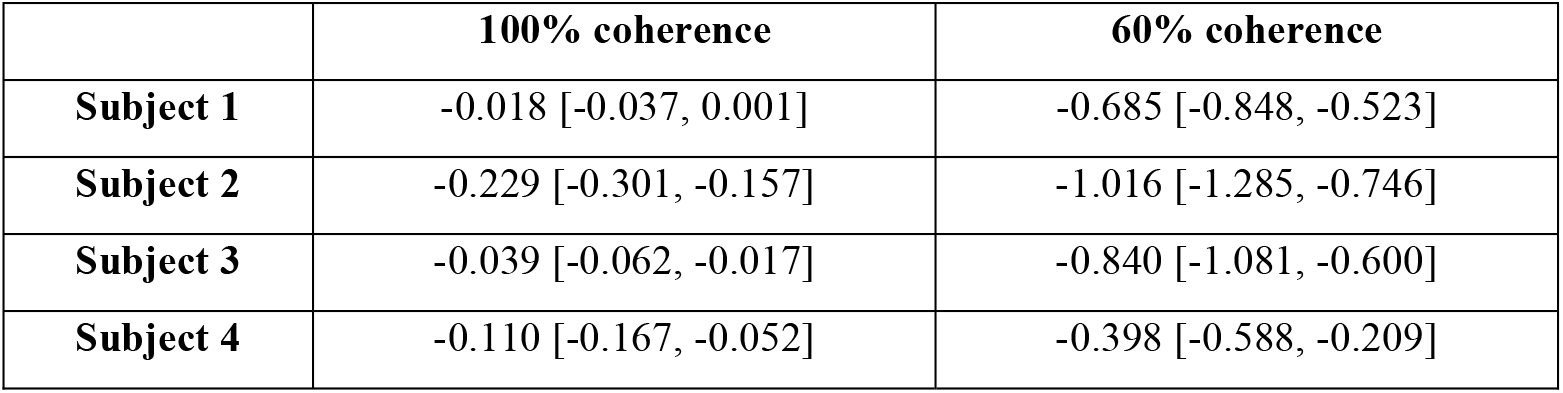
Individual-subject coefficients for target-condition × log_10_(background-speed) interaction. Linear models were fitted separately for each subject and coherence condition, with log10-transformed threshold ratio as the dependent variable and target condition, log10-transformed background speed, and their interaction as predictors. Values are the condition × log-speed interaction coefficients with 95% confidence intervals in brackets. The coefficient expresses the change in the decrement-increment difference in log_10_ threshold ratio per decade increase in background speed. Negative values indicate that the increment-decrement asymmetry decreases as background speed increases.

**STable 4.** Conservative group-level analyses across the four subjects. Each subject contributed a single summary value for each effect, and these values were tested against zero using two-tailed one-sample t tests (df = 3). Effects involving both coherence levels were calculated over the balanced 5-30°/s background-speed range. The target-condition effects are differences in mean log_10_-transformed threshold ratio between decrement and increment conditions. The coherence effect is the difference between 60% and 100% coherence, and the condition × coherence effect is the difference in the target-condition effect between 60% and 100% coherence. Speed-dependent effects were quantified as slopes against log_10_-transformed background speed. For the condition × speed effects, slopes were estimated over all available speeds within each coherence condition (2.5-30°/s for 100% coherence and 5-30°/s for 60% coherence). The coherence × speed effect is the slope of the coherence effect, averaged across target conditions, over the common 5-30°/s range; a negative value indicates that the effect of reduced coherence decreased with increasing background speed. For the condition × coherence × speed interaction, condition × speed slopes were estimated separately for the two coherence conditions over the common 5-30°/s range, and the 100%-coherence slope was subtracted from the 60%-coherence slope. A negative value therefore indicates that the increment-decrement asymmetry decreased more rapidly with increasing background speed under 60% than under 100% coherence.

| **Effect** | **Mean** | **t (df = 3)** | ***p*** |
|---|---|---|---|
| **Condition**<br>**100% coherence** | 0.059 | 3.32 | 0.045 * |
| **Condition**<br>**60% coherence** | 0.217 | 9.78 | 0.002 ** |
| **Coherence** | 0.164 | 11.30 | 0.002 ** |
| **Condition × Coherence** | 0.158 | 6.04 | 0.009 ** |
| **Condition × Speed**<br>**100% coherence** | -0.099 | -2.08 | 0.13 |
| **Condition × Speed**<br>**60% coherence** | -0.735 | -5.61 | 0.011 * |
| **Coherence × Speed** | -0.518 | -16.99 | < 0.001 *** |
| **Condition × Coherence × Speed** | -0.563 | -4.68 | 0.018 * |

**STable 5.** Weber-fraction regression slopes for individual subjects. For each subject and condition, log_10_-transformed Weber fraction was regressed on log_10_-transformed background speed, using individual staircase runs as observations. Values are the regression slopes with 95% confidence intervals in brackets. A slope of 0 is consistent with Weber’s law, in which the Weber fraction is constant across speeds; a negative slope indicates sub-Weber scaling, in which relative sensitivity improves as background speed increases. The 100% increment condition was fitted over 10–30°/s because the Weber fraction reached a local minimum near 5°/s (tested separately; see Results and Figure 5A); the other conditions were fitted over the full range of available background speeds.

| **Condition** | **Fit speed range (°/s)** | **Subject 1** | **Subject 2** | **Subject 3** | **Subject 4** |
|---|---|---|---|---|---|
| **100% coherence<br>Increment** | 10 - 30 | 0.10<br>[-0.07, 0.28] | 0.07<br>[-0.16, 0.30] | -0.05<br>[-0.24, 0.13] | 0.16<br>[-0.17, 0.47] |
| **100% coherence<br>Decrement** | 2.5 - 30 | -0.06<br>[-0.13, 0.02] | -0.35<br>[-0.45, -0.25] | -0.13<br>[-0.20, -0.05] | -0.15<br>[-0.23, -0.07] |
| **60% coherence<br>Increment** | 2.5 - 30 | -0.68<br>[-0.78, -0.58] | -0.46<br>[-0.60, -0.33] | -0.71<br>[-0.84, -0.58] | -0.88<br>[-0.98, -0.79] |
| **60% coherence<br>Decrement** | 5 - 30 | -0.78<br>[-0.88, -0.68] | -0.75<br>[-0.83, -0.67] | -0.64<br>[-0.77, -0.51] | -0.47<br>[-0.56, -0.39] |

## References

Albright, T. D. (1984). Direction and orientation selectivity of neurons in visual area MT of the macaque. Journal of Neurophysiology, 52(6), 1106–1130.

Albright, T. D., & Desimone, R. (1987). Local precision of visuotopic organization in the middle temporal area (MT) of the macaque. Experimental Brain Research, 65(3), 582–592.

Allman, J., Miezin, F., & McGuinness, E. (1985). Direction- and velocity-specific responses from beyond the classical receptive field in the middle temporal visual area (MT). Perception, 14(2), 105–126.

Benjamini, Y., & Hochberg, Y. (1995). Controlling the false discovery rate: a practical and powerful approach to multiple testing. Journal of the Royal Statistical Society: Series B (Methodological), 57(1), 289–300.

Born, R. T. (2000). Center-surround interactions in the middle temporal visual area of the owl monkey. Journal of Neurophysiology, 84(5), 2658–2669.

Born, R. T., & Bradley, D. C. (2005). Structure and function of visual area MT. Annual Review of Neuroscience, 28(1), 157–189.

Born, R. T., & Tootell, R. B. (1992). Segregation of global and local motion processing in primate middle temporal visual area. Nature, 357(6378), 497–499.

Braddick, O. (1993). Segmentation versus integration in visual motion processing. Trends in Neurosciences, 16(7), 263–268.

Brainard, D. H. (1997). The psychophysics toolbox. Spatial Vision, 10(4), 433–436.

Chakrala, A. S., Cao, Y., & Huang, X. (2023, November 11–15). Neural representation of multiple speeds of spatially separated stimuli in cortical area MT [Poster presentation PSTR082.13]. Neuroscience 2023, Society for Neuroscience, Washington, DC, United States.

Chakrala, A. S., Xiao, J., & Huang, X. (2024). The role of binocular disparity and attention in the neural representation of multiple moving stimuli in the visual cortex. BioRxiv, 2023–06.

Chuang, J., Ausloos, E. C., Schwebach, C. A., & Huang, X. (2016). Integration of motion energy from overlapping random background noise increases perceived speed of coherently moving stimuli. Journal of Neurophysiology, 116(6), 2765–2776.

De Bruyn, B., & Orban, G. A. (1988). Human velocity and direction discrimination measured with random dot patterns. Vision Research, 28(12), 1323–1335.

DeAngelis, G. C., Cumming, B. G., & Newsome, W. T. (1998). Cortical area MT and the perception of stereoscopic depth. Nature, 394(6694), 677–680.

Friedman, C. T., Wang, M., Yerxa, T., Arseneau, B. A., Huang, X., & Cooper, E. A. (2025). Statistical regularities in natural scenes that support figure-ground segregation by neural populations. PLOS Computational Biology, 21(10), e1013573.

Hohnsbein, J., & Mateeff, S. (2002). Speed decrements are seen better than increments through small apertures. Experimental Brain Research, 142(3), 434–438.

Huang, X., Albright, T. D., & Stoner, G. R. (2007). Adaptive surround modulation in cortical area MT. Neuron, 53(5), 761–770.

Huang, X., Albright, T. D., & Stoner, G. R. (2008). Stimulus dependency and mechanisms of surround modulation in cortical area MT. The Journal of Neuroscience, 28(51), 13889–13906.

Huang, X., Ghimire, B., Chakrala, A. S., & Wiesner, S. (2026). Neural coding of multiple motion speeds in visual cortical area MT. eLife, 13, RP94835.

Ivry, R. B., & Cohen, A. (1992). Asymmetry in visual search for targets defined by differences in movement speed. Journal of Experimental Psychology: Human Perception and Performance, 18(4), 1045–1057.

Krekelberg, B., & van Wezel, R. J. (2013). Neural mechanisms of speed perception: transparent motion. Journal of Neurophysiology, 110(9), 2007–2018.

Levitt, H. (1971). Transformed up-down methods in psychoacoustics. The Journal of the Acoustical Society of America, 49(2B), 467–477.

Lisberger, S. G., & Movshon, J. A. (1999). Visual motion analysis for pursuit eye movements in area MT of macaque monkeys. Journal of Neuroscience, 19(6), 2224–2246.

Lu, Z. L., & Dosher, B. (2013). Visual psychophysics: From laboratory to theory. MIT Press.

Masson, G. S., Mestre, D. R., & Stone, L. S. (1999). Speed tuning of motion segmentation and discrimination. Vision Research, 39(26), 4297–4308.

Maunsell, J. H., & Van Essen, D. C. (1983). Functional properties of neurons in middle temporal visual area of the macaque monkey. I. Selectivity for stimulus direction, speed, and orientation. Journal of Neurophysiology, 49(5), 1127–1147.

Newsome, W. T., & Paré, E. B. (1988). A selective impairment of motion perception following lesions of the middle temporal visual area (MT). Journal of Neuroscience, 8(6), 2201–2211.

Nover, H., Anderson, C. H., & DeAngelis, G. C. (2005). A logarithmic, scale-invariant representation of speed in macaque middle temporal area accounts for speed discrimination performance. Journal of Neuroscience, 25(43), 10049–10060.

Orban, G. A., Van Calenbergh, F., De Bruyn, B., & Maes, H. (1985). Velocity discrimination in central and peripheral visual field. Journal of the Optical Society of America A, 2(11), 1836–1847.

Pelli, D. G. (1997). The VideoToolbox software for visual psychophysics: Transforming numbers into movies. Spatial Vision, 10(4), 437–442.

Qian, N., & Andersen, R. A. (1994). Transparent motion perception as detection of unbalanced motion signals. II. Physiology. The Journal of Neuroscience, 14(12), 7367–7380.

Raiguel, S., Van Hulle, M. M., Xiao, D. K., Marcar, V. L., & Orban, G. A. (1995). Shape and spatial distribution of receptive fields and antagonistic motion surrounds in the middle temporal area (V5) of the macaque. European Journal of Neuroscience, 7(10), 2064–2082.

Recanzone, G. H., Wurtz, R. H., & Schwarz, U. (1997). Responses of MT and MST neurons to one and two moving objects in the receptive field. Journal of Neurophysiology, 78(6), 2904–2915.

Regan, D., Giaschi, D., Sharpe, J. A., & Hong, X. H. (1992). Visual processing of motion-defined form: selective failure in patients with parietotemporal lesions. Journal of Neuroscience, 12(6), 2198–2210.

Rudolph, K., & Pasternak, T. (1999). Transient and permanent deficits in motion perception after lesions of cortical areas MT and MST in the macaque monkey. Cerebral Cortex, 9(1), 90–100.

Salzman, C. D., Britten, K. H., & Newsome, W. T. (1990). Cortical microstimulation influences perceptual judgements of motion direction. Nature, 346(6280), 174–177.

Salzman, C. D., Murasugi, C. M., Britten, K. H., & Newsome, W. T. (1992). Microstimulation in visual area MT: effects on direction discrimination performance. The Journal of Neuroscience, 12(6), 2331–2355.

Sekuler, A. B. (1990). Motion segregation from speed differences: Evidence for nonlinear processing. Vision Research, 30(5), 785–795.

Snowden, R. J., Treue, S., Erickson, R. G., & Andersen, R. A. (1991). The response of area MT and V1 neurons to transparent motion. The Journal of Neuroscience, 11(9), 2768–2785.

Stocker, A. A., & Simoncelli, E. P. (2006). Noise characteristics and prior expectations in human visual speed perception. Nature Neuroscience, 9(4), 578–585.

Tadin, D., Lappin, J. S., Gilroy, L. A., & Blake, R. (2003). Perceptual consequences of centre– surround antagonism in visual motion processing. Nature, 424(6946), 312–315.

Wei, L., Mitchell, A. O., & Maunsell, J. H. (2023). Increments in visual motion coherence are more readily detected than decrements. Journal of Vision, 23(5), 18.

Weiss, Y., Simoncelli, E. P., & Adelson, E. H. (2002). Motion illusions as optimal percepts. Nature Neuroscience, 5(6), 598–604.

Whittle, P. (1986). Increments and decrements: luminance discrimination. Vision Research, 26(10), 1677–1691.

Wiesner, S., Baumgart, I. W., & Huang, X. (2020). Spatial arrangement drastically changes the neural representation of multiple visual stimuli that compete in more than one feature domain. The Journal of Neuroscience, 40(9), 1834–1848.

Wiesner, S., Ghimire, B., & Huang, X. (2026). Spatial preference-weighted representation of multiple stimuli in cortical area MT. The Journal of Neuroscience, 46(34), e1947252026.

Xiao, J., & Huang, X. (2015). Distributed and dynamic neural encoding of multiple motion directions of transparently moving stimuli in cortical area MT. The Journal of Neuroscience, 35(49), 16180–16198.

Zhang, L. Q., & Stocker, A. A. (2022). Prior expectations in visual speed perception predict encoding characteristics of neurons in area MT. The Journal of Neuroscience, 42(14), 2951–2962.

